# PEPerMINT: Peptide Abundance Imputation in Mass Spectrometry-based Proteomics using Graph Neural Networks

**DOI:** 10.1101/2024.03.23.586248

**Authors:** Tobias Pietz, Sukrit Gupta, Christoph N. Schlaffner, Saima Ahmed, Hanno Steen, Bernhard Y. Renard, Katharina Baum

## Abstract

**Motivation:** Accurate quantitative information about the protein abundance is crucial for understanding a biological system and its dynamics. Protein abundance is commonly estimated using label-free, bottom-up mass spectrometry protocols. Here, proteins are digested into peptides before quantification via mass spectrometry. However, missing peptide abundance values, which can make up more than 50% of all abundance values, are a common issue. They result in missing protein abundance values, which then hinder accurate and reliable downstream analyses.

**Results:** To impute missing abundance values, we propose PEPerMINT, a graph neural network model working directly on the peptide level that flexibly takes both peptide-to-protein relationships in a graph format as well as amino acid sequence information into account. We benchmark our method against eleven common imputation methods on six diverse datasets, including cell lines, tissue, and plasma samples. We observe that PEPerMINT consistently outperforms other imputation methods. Its prediction performance remains high for varying degrees of missingness, different evaluation approaches and differential expression prediction. As an additional novel feature, PEPerMINT provides meaningful uncertainty estimates and allows for tailoring imputation to the user’s needs based on the reliability of imputed values.

**Availability and implementation:** The code is available at https://github.com/DILiS-lab/pepermint.

## Introduction

Proteins are the main acting molecules in cells. The characterization of their quantity in different biological contexts plays a fundamental role in understanding cellular function and regulation in disease [1, 2]. Methods based on label-free mass spectrometry (MS) are commonly used for high throughput quantification of protein abundance in biological samples [3]. In MS-based bottom-up proteomics, proteins are enzymatically digested into peptides before subjecting them to a mass spectrometer. Individual peptides are then commonly identified by matching their spectra to corresponding databases [4]. With data-dependent acquisition (DDA), only the top most abundant peptides within a given analysis time window are individually fragmented and used for identification and quantification. In contrast, data-independent acquisition (DIA) fragments all peptides within a given time and mass window. The higher sensitivity of DIA has increased its use in recent years [5]. Finally, several aggregation methods exist to infer protein abundance by computationally aggregating the measured peptide abundance values into protein abundances [6, 7] to allow downstream analysis on the protein level.

With label-free MS, peptide abundance measurements exhibit a high number of missing values (e.g. 22.1% - 68.8% for the datasets used in this paper). These might either be due to peptides with an abundance below the detection limit, often referred to as missing not at random (MNAR), or due to random errors and stochastic fluctuations in the measurement process, often referred to as missing completely at random (MCAR) [8, 9, 10]. While performing peptide-to-protein aggregation, these missing values can propagate to the protein level and ultimately hamper downstream analyses [11, 12]. Therefore, different methods for imputing missing values following different paradigms - relying on single (e.g., minimal) values, leveraging local similarities or global structure - have been suggested and benchmarked [9, 12, 13] (see overview in Table 1). Basic methods, such as average, k-nearest neighbors (KNN), iterative singular value decomposition (ISVD), principal component analysis (PCA) or random forest (RF), that are applicable beyond proteomics have been especially widely adopted [14, 15]. More complex extensions, mostly dedicated to protein imputation, based on mixture models or matrix factorization, have been suggested [13, 16]. In addition, adaptations of basic methods such as RF or linear regression models have been proposed to include additional features like mRNA measurements [17].

**Table 1.** Overview of imputation methods used for our benchmark. We capture basic methods, more complex ones, and imputation methods based on deep neural networks representing all three generally considered categories of imputation methods. Further selection criteria were their appearance in proteomics imputation benchmark studies and their availability in terms of support in open-source software packages.

| Method | Benchmarked by | Supported by |
| --- | --- | --- |
|  | Single Value |  |
| MinDet | [9, 26] | [26, 27, 28, 29] |
| MinProb | [9, 12, 26, 30] | [26, 27, 28, 29] |
| Median | [19, 26] | [26] |
| Local Similarity |  |  |
| KNN | [9, 12, 15, 19, 26, 30] | [26, 27, 28, 29, 31] |
| RF [32] | [19, 26, 30] | [26, 27, 28, 29] |
| MICE [33] | [19, 26] | [26, 31, 33] |
| Global Structure |  |  |
| ISVD [34] | [9, 12, 26, 30] | [26, 35] |
| BPCA [36] | [12, 15, 26, 30, 19] | [26, 27, 28, 29, 35] |
| DAE [19] | [19] | [19] |
| VAE [19] | [19] | [19] |
| CF [19] | [19] | [19] |

Deep learning (DL) was suggested for missing value imputation in omics datasets. Arisdakessian et al. [18] introduced a basic neural network of several fully connected layers and a single dropout layer for imputing single-cell RNA sequencing datasets. Webel et al. [19] proposed the application of autoencoders for the imputation of MS-based proteomics datasets. Barzine et al. [20] used a neural network and mRNA expression values along with context information from GO terms and UniProt keywords to predict missing protein abundance values.

While additional information, such as mRNA measurements, can improve imputation performance, obtaining mRNA data requires costly additional wet-lab experiments or might even be infeasible (e.g., for plasma samples). Furthermore, there are multiple proteomics-specific features beyond GO terms or UniProt keywords that can provide additional information and context to machine learning models for learning patterns across similar proteins or peptides that current imputation methods fail to exploit. In particular, similarities in physical properties of the measured molecules, such as peptide mass, sequence length, and charge state, are helpful for peptide-to-protein aggregation [7]. Also, amino acid sequence information is available for missing proteins and peptides, and language models pre-trained on amino acid sequences have shown good performance on a variety of protein-related tasks [21]. The embeddings derived from these pre-trained language models encode the valuable biophysical properties of the underlying protein or peptide but currently remain unused as features for imputation. In addition, neither of the described DL-based imputation methods considers the particular relationships between proteins and peptides. Peptides originating from the same protein are expected to have strongly correlated abundances, a relationship that can be exploited to improve imputation performance and that also enables leveraging abundance information on non-unique peptides. Moreover, while multiple contributions have focused on imputing values in high missingness scenarios [16, 20], little attention has been paid to the inherent uncertainty coming with such imputations. So far, most imputation methods have not been designed with uncertainty in mind, resulting in uncertainty estimates for imputed values either being not available or obtained via multiple imputation [22]. Nevertheless, uncertainty estimates are of high value as they can enhance the trust in imputation results, and also help users filter out uncertain imputations.

We here address these gaps and propose a new DL-based model for imputation in proteomics datasets that exploits additional proteomics features in the form of amino acid sequences and peptide-protein relationships. As graph neural network (GNN) models have shown considerable success in modeling complex relationships between molecules and learning from biological and omics data [23, 24, 25], our method relies on a GNN architecture. What is more, while most proteomics imputation methods still impute on the protein level, our model acts directly on the peptide level, a strategy shown to yield improved imputation results [9]. Furthermore, our DL architecture enables uncertainty estimates for imputed values at low computational overhead to provide the user with a valuable tool for imputation prediction diagnostics. We systematically benchmark our novel method against eleven imputation methods from different categories across six representative datasets with different ground-truth mechanisms using three evaluation metrics (see overview in Fig. 1). Furthermore, we showcase its uncertainty quantification capabilities.

**Fig. 1.**
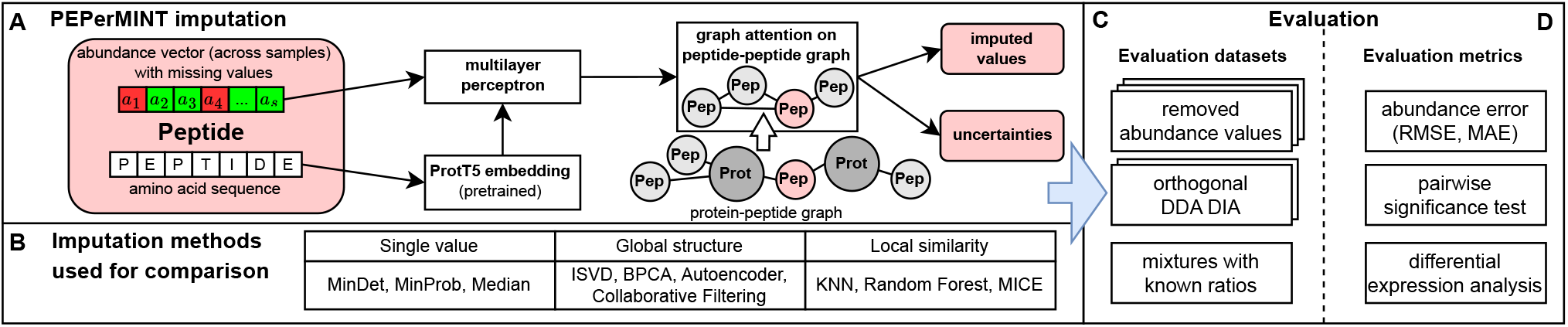
Overview of our PEPerMINT imputation method and our benchmarking framework. (A) Our PEPerMINT imputation model combines both peptide sequence information and abundance values across samples into a latent representation. Structural information is included via a peptide-peptide graph using a graph attention layer. (B) PEPerMINT is compared to eleven published imputation methods from three different categories. (C, D) We perform a systematic evaluation on six diverse datasets with ground truth derived from three different mechanisms with respect to three different evaluation metrics (see Methods for details).

## Methods

We introduce PEPerMINT (PEPtide Mass spectrometry Imputation NeTwork), a method combining abundance values and information from amino acid sequences and protein-peptide relations to impute missing values on the peptide level. For its implementation and systematic benchmarking, we use our novel open-source PyProteoNet framework (see Supplement).

### PEPerMINT imputation

For our PEPerMINT imputation model, we propose a neural network architecture combining a learnable transformation of abundance values, a GNN operating on the peptide graph, as well as amino acid sequence embeddings derived from a transformer-based language model (see Fig. 1A for a visual overview).

#### Input features

We assume a proteomics dataset with abundance values for *n* (potentially non-unique) peptides measured across *s* samples given as *n* × *s* matrix **A** where the elements of **A** either represent logarithmized (natural logarithm) and standardized (zero mean, unit variance) abundance values or missing values. Missing values are ignored for logarithmization and standardization. We address the problem of predicting abundance values for the missing values. PEPerMINT takes two inputs: the abundance matrix **A** and an *n* × 1024 sequence embedding matrix **S. S** is precomputed from the peptide amino acid sequences using the ProtT5 language model, which has previously shown good performance generating protein embeddings from sequence strings for tasks like predicting protein secondary structure [21]. This allows PEPerMINT to account for abundance values from non-missing samples as well as different biophysical peptide properties encoded in the sequence embeddings [21].

#### Peptide-peptide graph

The digestion of proteins into peptides for MS-based quantification results in the characteristic protein-peptide structure of MS-based datasets that can be described by a bipartite graph [42] where each peptide is assigned to one or more proteins. This structure can provide valuable information for the imputation of missing values since peptides belonging to the same protein are expected to show similar abundance profiles across samples. We transform this structure into a peptide-only graph *G* = (*V, E*) whereby peptides are nodes ∈ *V* that have an edge ∈ *E* between them if they belong to the same protein. Therefore, in *G*, all peptides belonging to the same protein are fully connected, and all peptides from proteins with shared peptides form a connected component (see Fig. 1A middle). We provide *G* as input to PEPerMINT.

#### Neural network architecture

Fig. 2 shows a simplified representation of PEPerMINT’s architecture. PEPerMINT scales down the sequence embeddings of each peptide by applying a learnable transformation 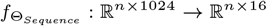. This aims to balance the size of abundance and sequence-based information. Next, for each peptide, we concatenate the sequence embedding and the vector containing peptide abundances across samples (abundance vector) and apply another learnable non-linear transformation to create a latent representation 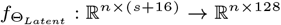.

**Fig. 2.**
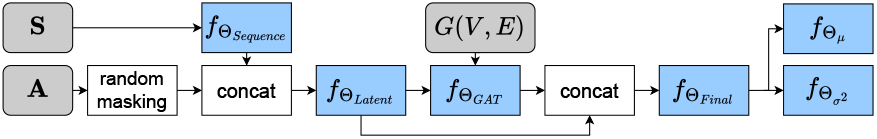
Simplified representation of the architecture of PEPerMINT with input feature representations (grey) and learnable (multilayer) transformations (blue). See Supplement Fig. S1 for a detailed visualization.

To account for the protein-peptide structure of the dataset represented by the peptide-peptide graph *G* we use an attention-based GNN consisting of a single GATv2 [43] layer with 64 heads with each head outputting a vector of shape 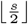. To keep the peptide-specific information from our latent representation, we add a skip connection bypassing the GNN. We add another learnable transformation on the concatenated output of the skip connection and the GNN output 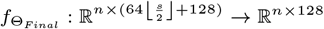

#### Uncertainty prediction of imputed values

To allow the estimation of uncertainty for imputed values, abundance values are predicted in a Bayesian setting. At the same time, this allows our model to better adapt to differing amounts of measurement noise for individual peptides (heteroscedastic noise) [44, 45]. Therefore, instead of single abundance values, mean and variance values of Gaussian abundance distributions are predicted [46] by two separate output heads [47] (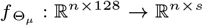 and 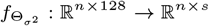).

#### Training scheme and self-supervised learning

We create a test set for each dataset by masking 10% of its abundance values uniformly at random (setting them to missing). However, for DDA/DIA datasets, the test set is given by all missing DDA abundance values that have a corresponding non-missing DIA value. From the remainder of the peptides (after picking the test set), we pick 10% of non-missing values uniformly at random as the validation set and mask them. On the resulting dataset, training is performed in a self-supervised manner. Similar to the training of denoising autoencoders as, e.g., done by Webel et al. [19], for each training step, we mask a fraction *γ* of non-missing values and compute the loss over them. The fraction *γ* is sampled randomly with samples uniformly distributed over the [5%, 15%) interval to improve model generalization.

To improve the training performance in the Bayesian setting, the model is trained in two rounds. First, we only train the 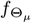 head (see Fig. 2) with mean squared error (MSE) loss before tuning the mean *µ* and variance *σ*^2^ together within a second training run using both output heads 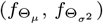 and Gaussian negative log-likelihood loss [48]. For both training rounds, we employ early stopping with respect to the MSE computed on the holdout validation set after each epoch. We define one epoch as consisting of 500 randomly masked datasets.

### Imputation methods used for comparison

To evaluate PEPerMINT, we compare it against a broad, representative selection of eleven methods from the literature that are commonly used for imputation and have appeared frequently in other proteomics benchmarks (see Table 1 and Supplement for details).

i. *Single-value methods*: These methods either impute missing abundance values with the same single value or, for each missing value, randomly draw a value from a predetermined distribution. We evaluated MinDet (using the 0.01 quantile of non-missing values within each sample), MinProb (drawing from a normal distribution around the 0.01 quantile within each sample), and Median (peptide-wise across samples) as commonly used methods of this class.
ii. *Local similarity methods*: These methods assume that missing values of a peptide can be predicted from the abundance values of similar peptides. We selected *k* nearest neighbor (KNN) imputation as it is simple and widely used [9, 26, 15]. We also included imputation based on a Bayesian ridge regression model as suggested by the MICE [33] imputation framework and an RF-based imputation [32] as a commonly used method with good performance reported previously [30].
iii. *Global structure methods*: These methods assume that proteomics datasets contain redundant information and can, thus, be well described by a low dimensional representation, which is leveraged for inferring missing values. We use Bayesian principal component analysis (BPCA) [14] and iterative singular value decomposition (ISVD) [34] as the most frequently used representatives. To include DL methods, two autoencoder (AE) based methods (variational AE and denoising AE) and a method based on collaborative filtering (CF), all recently proposed in [19], were considered.

### Datasets

We use six benchmark datasets with the goal of spanning a variety of biological backgrounds, varying degrees of complexity (blood plasma, cell lines, tumor tissue) with diverse dataset sizes (between *<*500 to *>*13000 proteins and *<*2600 to *>*100000 peptides) and differing percentages of missingness on the peptide level (22.8-68.1%) for evaluation with respect to three different types of ground truth (see overview in Table 2, and further details in the Supplement).

**Table 2.** Overview of benchmark datasets and their characteristics including availability via ProteomeXchange, dataset ground-truth category (masked: removed abundance values, DDA/DIA, mixture: mixture of known ratios), number of samples, number of biological samples (BS), technical replicates per biological sample (TR per BS), number of proteins and peptides, percentage of missing values on the peptide level.

| Name, Reference | Identifier | Category | #Samples | #BS | #TR per BS | #Peptides | #Proteins | Missingness |
| --- | --- | --- | --- | --- | --- | --- | --- | --- |
| A1: prostate cancer [37] | PXD029525 | masked | 18 | 6 | 3 | 57770 | 6292 | 54.0% |
| A2: Crohn’s fibrosis [38] | PXD022214 | masked | 13 | 13 | 1 | 37158 | 4481 | 33.7% |
| A3: breast cancer [39] | PXD035857 | masked | 15 | 15 | 1 | 103608 | 13627 | 68.8% |
| B1: HEK293-*E.coli* [40] | PXD018408 | DDA/DIA | 16 | 2 | 8 | 16400 | 3045 | 25.2% |
| B2: HIV blood [41] | PXD047528 | DDA/DIA | 15 | 15 | 1 | 2535 | 435 | 41.8% |
| C: HeLa-*E.coli* [6] | PXD000279 | masked + mixture | 6 | 2 | 3 | 50260 | 6683 | 22.1% |

The first three benchmark datasets (A1-A3) do not contain explicit ground truth values. Therefore, we mask abundances, as commonly done in the literature [9, 30], using the measured abundance of masked values as ground truth.

In addition, we use two datasets (B1-B2) acquired in DDA mode with orthogonal ground truth acquired in DIA mode. The more accurate DIA measurements contain fewer missing values, which allows the evaluation of imputation methods on genuinely missing values in the DDA data. To make the DIA and DDA data comparable, all DIA abundance values are scaled to have the same mean as the corresponding DDA abundance values.

For the evaluation of differential expression (DE), we use a dataset (labeled C) of protein mixtures with known (spiked-in) ratios from different organisms serving as ground truth. Similar datasets have been used in the literature to evaluate methods for peptide-to-protein aggregation [6] and imputation [15, 30].

### Evaluation metrics

For abundance-based evaluation, we use the root mean squared error (RMSE) on all missing values that have non-missing ground truth values (masked values or values with orthogonal DIA measurements) similar to earlier evaluations of imputation methods [15, 20, 30]. To allow variance estimation, we compute the RMSE sample-wise. As an additional abundance-based evaluation, we compare imputation methods with pairwise significance tests using a Bonferroni-corrected one-sided (paired) Wilcoxon signed-rank test. For every pair of imputation methods, the test compares the two absolute errors of imputed values for each peptide and dataset sample.

In addition, we evaluate imputation methods for the correct identification of differentially expressed peptides. For each peptide, the corresponding sample abundance values between groups of replicates of biological samples with different spike-in ratios are compared using a Benjamini-Hochberg corrected Welch’s t-test. Depending on the significance threshold, different peptides are detected as differentially expressed. Those are compared to a known ground truth of differentially expressed peptides (all spiked-in peptides in our mixture dataset C) to compute true positives and false positives. We assess the performance over varying significance thresholds via a ROC curve (see more details in Supplement section D).

## Results

We evaluated the performance of our PEPerMINT peptide imputation methods on six proteomics datasets with various biological backgrounds and missingness characteristics. Then, we also compared it against a broad, representative selection of eleven widely used imputation methods. A comparison of the runtime of all presented imputation methods can be found in Fig. S8 of the Supplement.

### Abundance-based evaluation

We first performed an abundance-based evaluation via the sample-wise RMSE for four datasets with artificially introduced missing values and two DDA/DIA datasets with ground truth values acquired using the DIA (Fig. 3A). Particularly, we find that our PEPerMINT imputation method gives the best performance across all evaluated datasets, outperforming the second best-performing method (BPCA) by up to 20% on the breast cancer dataset. Out of the other evaluated methods, RF, BPCA, MICE, and CF also show good results. Interestingly, imputing missing values with the peptide-wise median gives better results than the more complex KNN imputation and methods based on autoencoders (DAE, VAE). ISVD, MinDET, and MinProb imputation were found to be generally worse, except for the good performance of MinDet and MinProb on the HIV blood dataset. We obtain similar results for dataset-wise mean absolute error as an alternative metric (see Supplement).

**Fig. 3.**
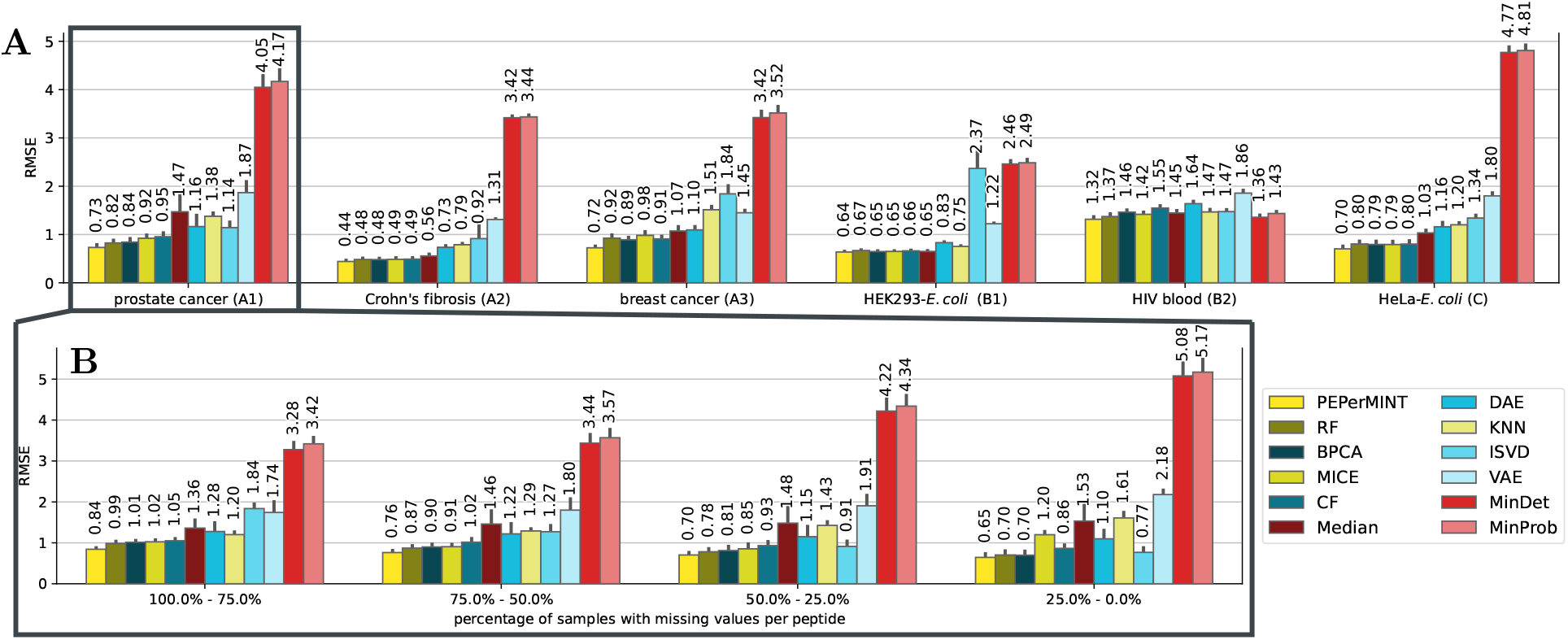
A: Sample-wise RMSE of all evaluated imputation methods on all six benchmark datasets with 95% confidence interval error bars (bootstrapped). B: Results for the prostate cancer (A1) dataset stratified by their fraction of missing values over the samples (see supplement for stratified results of other datasets). Our newly proposed PEPerMINT imputation outperforms other methods on all datasets, irrespective of the missingness fraction.

Further, as predicting the missing abundance of a peptide could be hampered if measured only in a few samples, we investigated whether imputation performance depends on the degree of missingness per peptide. Therefore, we stratified the evaluated peptides by their fraction of missing values across samples (Fig. 3B for the prostate cancer dataset; similar results for other datasets, see Supplement). Again, PEPerMINT outperforms other imputation methods for any fraction of missing values. The biggest advantage over competitor methods is observed on peptides with high fractions of missing values. It can also be noted that PEPerMINT, RF, BPCA, CF, DAE, and especially ISVD imputation show improved performance when the fraction of missing values decreases. In contrast, MinProb and MinDet perform better with high fractions of missing values.

Fig. 4 shows the results for statistically comparing imputation methods using a Wilcoxon signed-rank test (see Methods). Our PEPerMINT method performs significantly better than all other evaluated methods on the majority of benchmark datasets. It should be noted that in contrast to the sample-wise RMSE results shown in Fig. 3, the Wilcoxon test compares individual imputed values without averaging the error per dataset sample. The good performance of PEPerMINT also holds when stratifying peptides for missingness across samples (see Supplement Fig. S4).

**Fig. 4.**
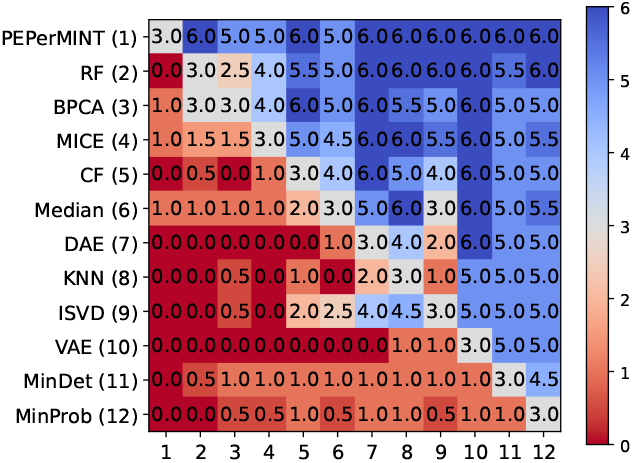
Pairwise comparison of different imputation methods by statistical significance test results (see Methods). Colors encode how many of the six evaluated datasets the imputation method given in the row performs significantly better (5% significance level) than the imputation method given in the column (insignificant test results increase the count by 0.5). Blue cells indicate the method given in the row outperforms the imputation method given by the column in the majority of cases.

### Evaluation of differential expression prediction

DE analysis is a common downstream analysis task performed on MS-based proteomics datasets. Therefore, we compared our proposed method with the other imputation methods with respect to the performance of DE analysis on the imputed dataset. For evaluation, we used the ground truth protein ratios that can be inferred from the species-specific mixture rates. We restricted the evaluation to peptides that can uniquely be assigned to one species. The receiver operating characteristics (ROC) curve in Fig. 5 shows that our method is performing better than the other methods, with the highest area under the ROC curve (AUC). The precision-recall curve (Supplement Fig. S5) supports this result.

**Fig. 5.**
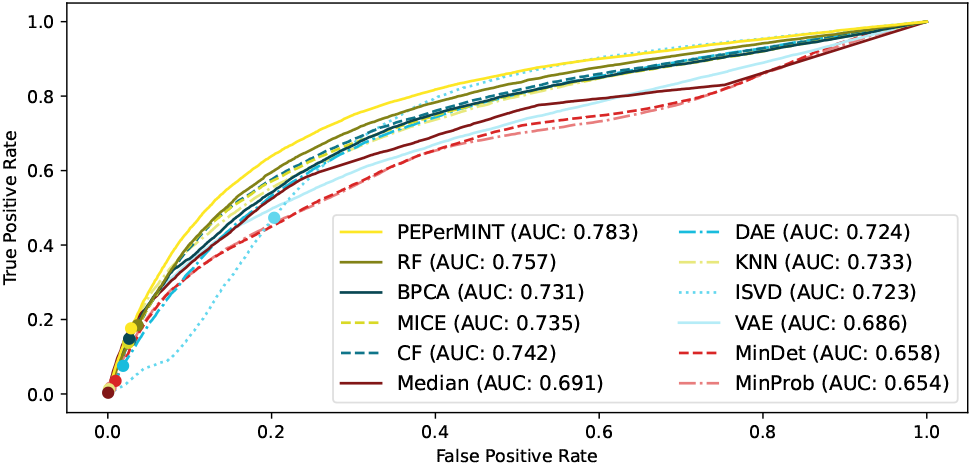
ROC curve for the performance of DE analysis of peptides on the HeLa-*E*.*coli* dataset imputed with different imputation methods with 5% FDR thresholds marked (dots). Our PEPerMINT imputation method (yellow) outperforms other methods, having the largest area under the curve (AUC).

### Predicted uncertainty of imputed values

Our PEPerMINT approach allows the out-of-the-box prediction of uncertainty for imputed values, helping users obtain a quantitative estimate of their trustworthiness. To evaluate the usefulness of this computed uncertainty, we compared the imputed values against their ground truth colored by their predicted uncertainty (Fig. 6A). The imputed values with the lowest uncertainty (dark blue) tend to show better predictions (low error) and high abundances. The latter fits with the characteristics of data acquired via MS because high abundance values commonly are proportionally less influenced by measurement noise and are, therefore, assumed to be more reliable [49]. In addition, we find that removing imputed values with high predicted uncertainty from the evaluation generally improves imputation quality (Fig. 6B). Furthermore, we observe that filtering out imputations with substantial uncertainty but keeping those with low uncertainty, can again massively increase the accuracy of downstream analysis. Reusing the experimental setup of the DE analysis in Fig. 5, we find that by filtering at a predicted uncertainty threshold of 0.2 in Fig. 6C, we can obtain an AUC of 0.84, compared to an AUC of 0.78 for the PEPerMINT imputation alone (vs. 0.68 without imputation, see Supplement). This further validates the quality and benefit of the uncertainty predictions given by our method.

**Fig. 6.**
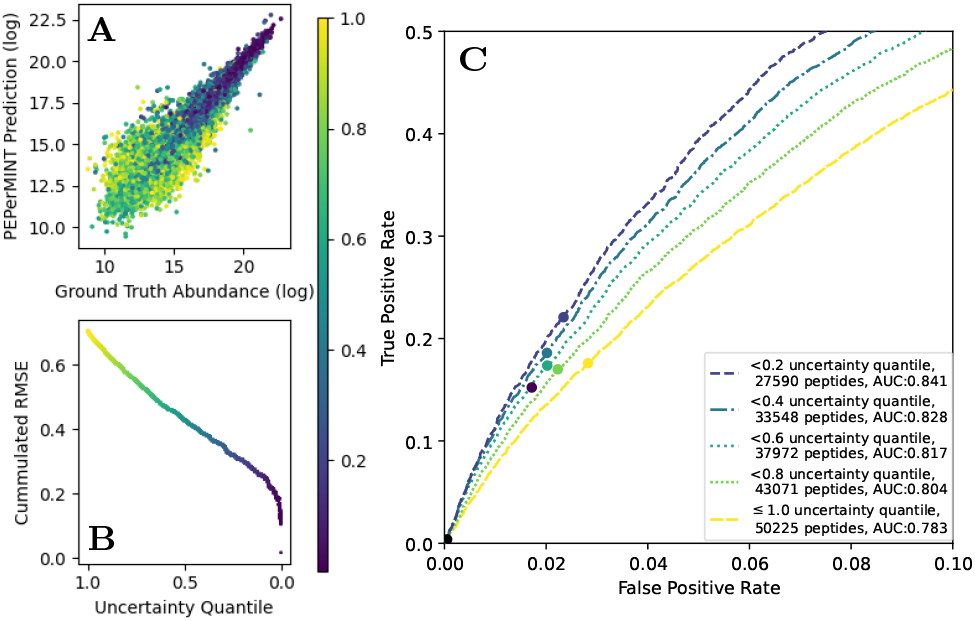
PEPerMINT’s predicted uncertainty for the imputed values for the HeLa-*E*.*coli* dataset (see Supplement for other datasets). A: Imputed abundance values vs. ground truth colored by predicted uncertainty (low: dark blue, high: yellow). B: Imputed values ordered by their predicted uncertainty with RMSE computed over different uncertainty quantiles [50]. C: ROC curve zoomed in at low FDR values for the performance of DE analysis for imputing values only up to a predicted uncertainty threshold (see Supplement for details). The yellow ROC curve (*≤* 1.0 uncertainty) is identical to the yellow PEPerMINT ROC curve from Fig. 5. Filtering out imputed values with high predicted uncertainties decisively improves DE analysis performance after imputation.

## Discussion and conclusion

Overall, PEPerMINT results in superior performance compared to other benchmarked methods across datasets, missingness levels of the peptides, and evaluation metrics. In addition, PEPerMINT provides a handle to the problem of imputation quality by predicting uncertainties for imputed values, with evident improvement potential for downstream analyses. This also distinguishes PEPerMINT from most other imputation methods, which commonly cannot result in confidence statements. We showed that PEPerMINT’s uncertainty estimates are highly correlated with imputation error, thereby aptly guiding users on when to rely on or filter out the imputed values. Further, filtering out imputed peptide abundance values with high predicted uncertainty eventually decisively improved the performance of the DE prediction task.

From the diverse characteristics of our benchmark datasets, it can be derived that PEPerMINT’s high performance is not limited to a specific dataset size or fraction of missing values. Further, our benchmark comprises datasets with and without technical replicates, i.e., samples with very high similarity. Thus, imputation could be considered easier when relying on technical replicates. However, PEPerMINT’s performance seems unaffected by this factor and even outperforms other methods by the largest margin on the breast cancer dataset, which is devoid of technical replicates. This further hints at PEPerMINT actually learning biologically relevant patterns instead of merely averaging across technical replicates.

Comparing our different categories of datasets, PEPerMINT performs best on our masked benchmark datasets that, by design, only exhibit MCAR missing values. This can be explained by our self-supervised training scheme, which also masks uniformly at random and aligns well with MCAR missing values. Nevertheless, PEPerMINT still shows very good results on DDA benchmark datasets with DIA ground truth values that can be assumed to contain both MCAR and MNAR missing values. Further, PEPerMINT also performs well on the HIV blood dataset, in which a high fraction of missing values is due to lowly abundant peptides (MNAR) as the blood plasma proteome is well studied with missing values rarely occurring. Its increased percentage of MNAR compared to the other datasets could be the cause for the different ranking of imputation method performances on this dataset, e.g., very good performance of MinDet and MinProb imputation that replace missing values with low abundance values.

For peptides with a high percentage of missing values, PEPerMINT compares especially well against other well-performing methods such as BPCA or RF imputation. This can be explained by PEPerMINT’s ability to exploit additional information (amino acid sequence, abundance of peptides belonging to the same protein) to obtain context about a peptide’s properties, even if little abundance information is available for the peptide itself. Indeed, using ablation studies, we find both additional information layers to provide at least some performance benefit to PEPerMINT (see Supplement Fig. S9). As our method also allows the flexible integration of other information layers both in tabular as well as graph form, it could be readily extended to improve proteomics imputation even further.

A limitation of our method when compared with other imputation methods is its higher runtime (see Supplement Fig. S8). However, the fastest-running methods like Median or MinProb also perform worse than more complex methods with longer runtimes like BPCA or RF. When executed on a GPU, PEPerMINT shows a runtime similar to or faster than that of BPCA imputation. Of note, all considered imputation methods finish within minutes, which is well acceptable for MS-based proteomics analysis workflows.

Further benchmarking criteria [51] and methods for proteomic imputation relying on DL and ensembling [52] or statistical models that take the protein-peptide structure into account [53] are emerging. They are exciting avenues for future exploration and for potential extensions of PEPerMINT, our GNN-based method working directly on the peptide level that flexibly takes both peptide-to-protein relationships as well as amino acid sequence information into account to improve prediction of missing abundance values.

## Data and Code Availability

All datasets, with the exception of the HIV blood dataset, were downloaded from the links provided in their original publications. The repository containing the code used for all experiments can be found under https://github.com/DILiS-lab/pepermint. The repository also contains links to relevant dataset files extracted from the original datasets (also including the HIV blood dataset).

## Competing interests

No competing interests are declared.

## Acknowledgements

This work received funding from the Klaus Tschira Foundation gGmbH (KTBoost KT25/GSO, to KB) and the Deutsche Forschungsgemeinschaft (DFG grant number RE2474/5-1, to BYR).

## Notes

### Competing Interest Statement

The authors have declared no competing interest.

