## Supplementary Information for "PEPerMINT: Peptide Abundance Imputation in Mass Spectrometry-based Proteomics using Graph Neural Networks"

**D Evaluation procedure for differential expression analysis** **8**

**E Implementation** **8**

**F Supplementary evaluation results** **9**

**G Runtime** **14**

**H Ablation study** **15**

### A PEPerMINT

Here, we show the detailed architecture of our proposed PEPerMINT deep neural network and state hyperparameters used for training.

#### A.1 Detailed neural network architecture

Figure S1 shows PEPerMINT's detailed architecture. The units highlighted in red correspond to the simplified architecture shown in Fig. 2 in the main text.

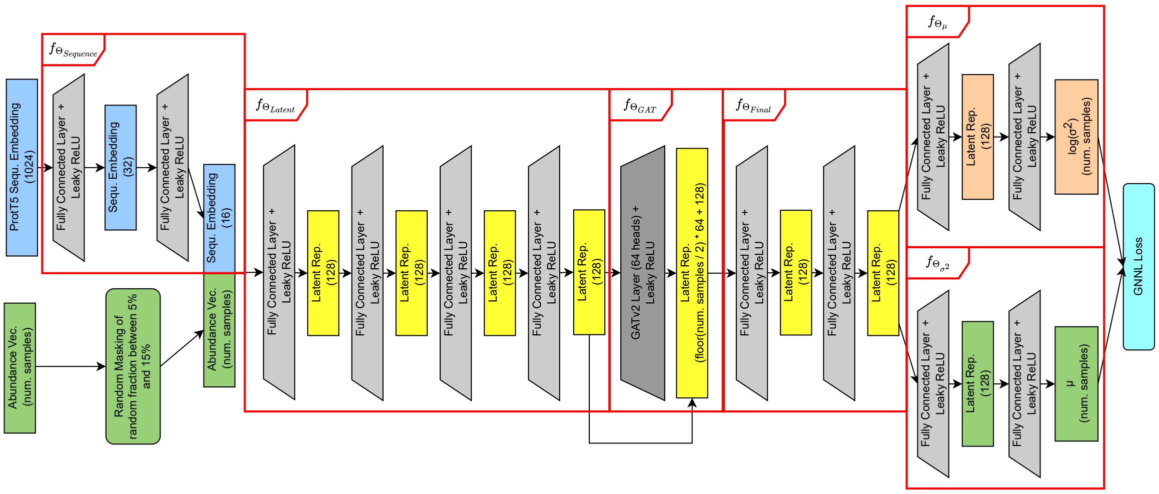

Fig. S1: Detailed architecture of PEPerMINT combining fully connected as well as graph attention layers. The units highlighted in red correspond to the components of the simplified architecture shown in Fig. 2 in the main text.

#### A.2 Training details

Before training, we ensure that all peptide abundance values that are part of the holdout test set (masked values for datasets A1, A2, A3, C; missing DDA values with DIA abundance values for B1, B2) are replaced by missing values (`NaN`). In addition, we create a holdout validation set by masking (replacing with `NaN`) 10% of the remaining non-missing values. Since our neural network requires all values to be numeric, we replace missing values (both masked and genuine missing) with a constant abundance value $c = -3$ (representing an abundance value smaller than 99.8 of all values, assuming a normal distribution of standardized values). We found that a missing constant $c$ representing a small abundance value works best for neural-network-based imputation.

On the resulting dataset, training is performed in a self-supervised manner. Similar to the training of denoising autoencoders as, e.g., done by Webel et al. [1], for each training step, we mask a fraction $\gamma$ of non-missing values and compute the loss over them. The fraction $\gamma$ is sampled randomly with samples uniformly distributed over the $[5\%, 15\%)$ interval to improve model generalization.

As described in the main text, the model is trained in two rounds to improve the training performance in the Bayesian setting. First, we only train the $f_{\Theta_\mu}$ head (see Fig. S1) with MSE loss before tuning the mean $\mu$ and variance $\sigma^2$ together within a second training run using both output heads ($f_{\Theta_\mu}$, $f_{\Theta_{\sigma^2}}$) and Gaussian negative log-likelihood (GNLL) loss as shown in equation 1 [2, 3, 4]. This loss function, for every masked value, computes the loss from the predicted abundance mean $\mu$, variance $\sigma^2$, and the original abundance value (before masking) $a$. The $\epsilon$ hyperparameter is set to 0.000001. The final loss is averaged across all abundance values that were masked for training. For both training rounds, we employ early stopping with respect to the MSE computed on the holdout validation set after each epoch. We set the early stopping patience to three epochs.

$$\mathcal{L}_{GNLL}(\mu, \sigma^2, a) = \frac{1}{2}\left(\log\left(\max\left(\sigma^2, \epsilon\right)\right) + \frac{(a-\mu)^2}{\max\left(\sigma^2, \epsilon\right)}\right) \tag{1}$$

For training, we use stochastic gradient descent with Adam as an optimizer and a learning rate of 0.0005. In addition, to make the training more memory efficient, we subsample the graph. Therefore, for each training step, we sample a subgraph induced by 50% of all peptide nodes together with their full neighborhood. We define one epoch as consisting of 500 randomly masked and subsampled graphs.

### B Imputation methods used for comparison

Here, we provide further details for the imputation methods from the literature that we compared against. All imputation methods are applied to logarithmic abundance values. It is further assumed that all logarithmic abundance values are represented by a matrix $\mathbf{A}$ with rows representing peptides and columns representing samples. For example, a dataset of $m$ peptides and $s$ samples is represented by an $m \times s$ matrix $\mathbf{A}$ with $\mathbf{A}_{i,j}$ containing the logarithmic abundance values of the $i$th peptide in the $j$th sample which can either be a real value or `NaN` in case of a missing value.

#### B.1 Single-Value methods

Single-value methods either impute multiple missing abundance values with the same single value or, for each missing value, randomly draw an imputed value from a predetermined distribution. Here, we consider three representative methods referred to as *MinDet*, *MinProb*, and *Median*.

##### MinDet

All missing values of one sample are replaced with a single value close to the minimum of the non-missing values within the sample. We use the 0.01 quantile of all non-missing abundance values within a sample.

##### MinProb

Missing values are replaced by random draws from a normal distribution centered around the 0.01 quantile of non-missing values per sample. The standard deviation of the normal distribution is estimated by the median of the row-wise standard deviations in $\mathbf{A}$, only considering rows with more than 50% of non-missing values. We use the implementation provided by the R package imputeLCMD [5].

##### Median

For every peptide, missing values are replaced with the median of all non-missing abundance values for this peptide in other samples.

#### B.2 Local similarity methods

Local similarity imputation methods assume that missing values can be predicted from the abundance values of a limited number of similar peptides. We consider KNN imputation as a simple and widely used local similarity method [6, 7, 8, 9]. In addition, imputation using random forests (RFs) is considered as a second representative local similarity method, which has been reported to show good performance [9, 10]. Finally, we include imputation based on a Bayesian ridge regression model as suggested by the commonly used MICE [11] imputation method.

##### K Nearest neighbors (KNN)

For a peptide $i$ with a missing value in sample $j$, the $k$ most similar proteins or peptides with non-missing abundance values for sample $j$ are identified. The similarity is determined by the Euclidean distance between the abundance profiles across samples [12] (only considering non-missing values). We use the implementation provided by the Scikit-learn Python package [13], $k$ is set to 5.

##### Random forest (RF)

For this imputation method, missing value imputation is formulated as a regression problem using a RF [14]. Therefore, all missing values are first replaced with an initial value. Then, for every column (sample) $j$ in $\mathbf{A}$, an RF is fit with the non-missing values for sample $j$ as response variables and all remaining non-missing abundance values of other samples as predictor variables. Imputation is then done by predicting missing values for column $j$ using the previously trained RF. This process is repeated for all samples and then iterated until a stopping criterion is met [14]. We use the missingpy[1] Python package.

##### MICE (Bayesian ridge regression)

With this method, missing values for one peptide within one sample are predicted from the abundance values for this peptide in other samples using a Bayesian ridge linear regression model. Imputation is done in an iterative fashion, where all missing values are first replaced with the per-sample mean. Subsequently, for each sample, a Bayesian ridge regression model is trained on the non-missing abundance values of the sample using the remaining samples as predictor variables. The trained model is then used to predict the missing values for the sample. This process is repeated for all samples and iterated until a convergence threshold is reached [11] We use the implementation of the IterativeImputer provided by the scikit-learn Python package [13], which is based on the original implementation by Van Buuren and Groothuis-Oudshoorn [11].

#### B.3 Global structure methods

Global structure imputation methods assume that proteomic datasets contain redundant information and can, therefore, be well described by a low dimensional representation, which can also be used to infer missing values. We use Bayesian principal component analysis (BPCA) [15] and iterative singular value decomposition (ISVD) [12] as two representative, frequently used methods. In addition, the two autoencoder-based methods recently proposed by Webel et al. [1] can also be categorized as global structure methods.

##### Iterative singular value decomposition (ISVD)

ISVD can be used for missing value imputation by decomposing $\mathbf{A}$ as shown in equation 2 where $\mathbf{U}$ and $\mathbf{V^T}$ are unitary matrices and $\mathbf{\Sigma}$ is a rectangular diagonal matrix. Columns of $V$ can be referred to as eigenpeptides with their corresponding squared eigenvalues given by $\mathbf{\Sigma}$. It is further assumed that every peptide can be represented by a linear combination of the $k$ eigenpeptides with the largest eigenvalues, with $k$ representing a tunable hyperparameter. Imputation is done by first replacing all missing values with a constant $c$ before decomposing $\mathbf{A}$. Missing abundance values for a peptide are then regressed as a linear combination of the $k$ eigenpeptides, where the regression coefficients are determined from the non-missing abundance values for this peptide. This process is then iterated until a convergence threshold is reached [16, 12].

$$\mathbf{A} = \mathbf{U\Sigma V}^T \tag{2}$$

We use the ISVD implementation provided by the fancyimpute Python package [17]. Following Troyanskaya et al. [12], the $k$ parameter is set to 20% of the overall number of samples rounded up to the next integer, and $c$ is set to 0.

[1] `https://github.com/epsilon-machine/missingpy`

##### Bayesian principal component analysis (BPCA)

Principal-component-based imputation methods like BPCA infer missing values from a linear combination of principal components. Those are calculated from the eigenvectors of the $s \times s$ covariance matrix $\mathbf{S}$ of $\mathbf{A}$.

Like ISVD imputation, principal-component-based imputation assumes that missing values can be regressed as a linear combination of $k$ principal axes inferred from the eigenvectors with the largest eigenvalue of $\mathbf{S}$. BPCA transfers this approach into a Bayesian framework. While a detailed description of the method can be found in the original paper by Oba et al. [15], one notable advantage of the Bayesian approach is the increased robustness with respect to the hyperparameter $k$. Since redundant principal axes are automatically suppressed, $k$ can always be set to its maximum value of $s - 1$, eliminating $k$ as a hyperparameter [15]. We use the implementation provided by the pcaMethods R package [18].

##### Autoencoder

Similar to ISVD and BPCA, autoencoder methods are built on the assumption that a high-dimensional input can be reconstructed from a low-dimensional representation. However, in contrast to principal-component-based methods, this low-dimensional latent representation is learned by an autoencoder neural network [19]. The corresponding network weights and, thereby, the latent representation are learned via stochastic gradient descent. For better performance and faster convergence during training, $\mathbf{A}$ is standardized to have zero mean and unit standard deviation [20]. In addition, to provide a valid input for the autoencoder neural network, missing values are replaced by the molecule-wise mean in the training set.

Using the implementation by Webel et al. [1], two types of four-layer autoencoders are used for imputation. For both setups, the input is given by the $m$ dimensional column vectors of $\mathbf{A}$ representing the peptide abundance values for single samples. Training is done on one sample at a time. The size of the latent representation is set to 50. In the differential autoencoder (DAE) setup, the autoencoder is prevented from learning the identity function by some of the abundance values being masked in the input. More specifically, 20% of the non-missing abundance values are chosen uniformly at random and replaced by 0 (representing the dataset abundance mean after normalization) [1]. Mean squared error (MSE) is used as a loss function, with the loss only being computed on originally non-missing abundance values. Dropout of 20% and batch normalization are used for the hidden layers. The alternative variational autoencoder (VAE) setup does not randomly mask the input vector but instead uses a probabilistic formulation of the latent vector as a multivariate Gaussian distribution. The loss function consists of a reconstruction loss combined with a regularization term regulating the standard normal distribution of the latent vector. For the exact loss function, it is referred to Webel et al. [1].

##### Collaborative filtering

In addition to autoencoders, collaborative filtering was recently suggested as an imputation method for proteomics datasets [1]. With this method, each peptide (row) and sample (column) is represented by a learnable embedding of fixed size $k$. Let $\mathbf{e}_i$ be the embedding for a peptide $i$ and $\mathbf{f}_j$ that for a sample $j$. Embedding values are learned from the non-missing values by minimizing the mean squared error (MSE) between the dot product between a peptide embedding and a sample embedding and the corresponding non-missing abundance value in $A$ as shown in equation 3. In this equation $non\_missing(\mathbf{A})$ refers to the function returning all indices of non-missing values for the matrix $\mathbf{A}$.

$$MSE_{CF}(\mathbf{A}) = \frac{\sum\limits_{i,j \in non\_missing(\mathbf{A})} ((\mathbf{e}_i \cdot \mathbf{s}_j) - \mathbf{A}_{i,j})^2}{|non\_missing(\mathbf{A})|} \tag{3}$$

We use the implementation provided by Webel et al. [1] with an embedding size of 15.

### C Datasets

Table S1 gives an overview of the mass spectrometry instruments as well as the analysis software and its parameters with respect to post-translational modifications used for the analysis of the raw files for our benchmark datasets. For more information, we refer to the original publication for each dataset.

| dataset | dataset identifier | instrument | analysis software | fixed modifications | variable modifications |
|---|---|---|---|---|---|
| prostate cancer | A1 | Q Exactive HF | MaxQuant | Carbamidomethyl(C) | Oxidation(M), Acetyl(N-term) |
| Crohn's fibrosis | A2 | Q Exactive Plus | MaxQuant | - | Oxidation(M) |
| breast cancer | A3 | Orbitrap Fusion Lumos | MaxQuant | - | Oxidation(M) Acetyl(N-term), Deamidation(NQ) |
| HEK293-*E.coli* | B1 | Thermo Scientific Q-Exactive | MaxQuant (DDA) Spectronaut (DIA) | Carbamidomethyl(C) | Oxidation(M), Acetyl(N-term) |
| HIV blood | B2 | timsTOF Pro 2 | FragPipe (DDA) Spectronaut (DIA) | Carbamidomethyl(C) | Oxidation(M) |
| HeLa-*E.coli* | C | LTQ Orbitrap | MaxQuant | Carbamidomethyl(C) | Oxidation(M), Acetyl(N-term) |

Table S1: Overview of instrument, software, and considered post-translational modifications used for the creation of the benchmark datasets.

#### C.1 Retrieval and LC/MS analysis of the HIV blood dataset

For the HIV blood dataset, we provide some further details on the generation and raw file processing for generating DDA and DIA abundance values.

##### C.1.1 Sample processing method

Our cost-effective method includes the addition of 3.5% (final concentration) perchloric acid, which results in protein precipitation and separation into a soluble and insoluble protein fraction, with the soluble plasma proteins being analyzed by LC/MS. The details about the method can be found [21].

##### C.1.2 LC/MS-based plasma proteome mapping

Samples were analyzed with a nano Elute liquid Chromatography (Bruker) coupled to a timsTOF Pro 2 mass spectrometer (Bruker Daltonics, Billerica, MA). Two microliters equivalent to 200ng of tryptic digest of the supernatant were loaded onto a C18 UHPLC column 50 mm x 150 $\mu$m from IonOpticks (Fitzroy, Australia). Peptides were separated by a 7 min gradient from 2% B to 34% B at a flow rate of 2 Ml/min (total run time $\approx$ 15 min). For the Data Dependent Analysis (DDA), the parameters were set as follows: m/z range 100–1700, the ion mobility (1/K0) range was set to 0.7–1.45 Vs/cm2, and the accumulation time was set to 100 ms.

##### C.1.3 LC/MS data analysis

All DDA raw files were searched with MSFragger software v3.1.1 [22] against a Human protein sequence without isoforms downloaded from UniProt, allowing for the following protein modifications: carbamidomethylated cysteine residues (fixed), oxidation of methionine (variable). The data were searched using the match between runs quantification parameter. All other parameters were set to default, including quantification by MaxLFQ.

The DIA runs were analyzed in Spectronaut version 13.12.200217 (Biognosys). Default BGS Factory settings were employed for both library generation with Pulsar and DIA analysis. Only DIA data were used for identification and quantification of proteins, and the subsequent analysis. Results were filtered with a 1% FDR at both the precursor and protein level.

#### C.2 Dataset preprocessing and evaluation

Here, we describe in more detail the generation and preprocessing of our datasets. To prevent data leakage, we made sure that no duplicated peptides (multiple peptides with the same amino acid and same sequence) were in the datasets. If there were duplicates (only the case for the HIV blood dataset and the DIA part of the HEK293-*E. coli* dataset), we aggregated all duplicates into one peptide by summing their abundance values. Afterward, preprocessing is done differently for each dataset category (A, B, C).

##### C.2.1 A: Masked datasets

Three different datasets are used: A1: This dataset (PXD029525) contains three prostate cancer cell cultures, each containing a treated and untreated condition group [23]. Each of the six biological samples (condition groups) consists of three technical replicates. A2: This dataset (PXD022214) consists of 13 tissue samples from five Crohn's disease patients with different grades of fibrosis [24]. No technical replicates are present A3: This dataset (PXD035857) contains 129 breast cancer patient-derived xenograft (PDX) tumors transplanted to mice [25] without technical replicates. Due to the increased runtime, especially for resource-intensive imputation methods like BPCA or RF (also noted by Webel et al. [1]), we keep computations manageable by randomly selecting 15 samples out of the original dataset.

We create each dataset from the peptides.txt file output of MaxQuant retrieved from the corresponding PRIDE repository entry. We use the "Intensity" columns in that file as our peptide abundance values. Protein-peptide relations are inferred from the "Protein group ID" column. The amino acid sequence of each peptide is inferred from the "Sequence" column. For the breast cancer dataset (A1), we subsample the dataset to 15 samples as described above. More specifically, the following 15 samples were randomly chosen: 33070, 33072, 33073, 33110, 33112, 33116, 33118, 33867, 38703, 38710, 38712, 38716, 38719, 38730, 39474. We create a hold-out test set for each dataset by masking. Therefore, we uniformly at random replace 10% of the non-missing peptide abundance values within each dataset with missing values ($NaN$). Afterward, we remove all peptides that show missing values across all samples.

##### C.2.2 B: DDA datasets with DIA ground truth

We use two DDA datasets with corresponding DIA ground truth. B1: This dataset (PXD018408) consists of human proteins (HEK293 cell lysate) with *E. coli* proteins spiked in at two different ratios [26] (1:1 and 1:2). All data is loaded from the data.xlsx file provided as supporting information by the original authors. DDA peptide abundance values are loaded from the LFQ intensity columns of the "Table S3 MaxQuant Output" table with amino acid sequence strings taken from the "Sequence" column and protein-peptide relations inferred from the indices given in the Protein group IDs column. DIA peptide abundance values are loaded from the "Table S4 Spectronaut Output" table with amino acid sequence strings extracted from the "EG.PrecursorID" column. Resulting duplicated peptides (same sequence) are grouped together by summing their abundance values. Matching between DDA and DIA peptides is done via amino acid sequence. B2: This clinical dataset (PXD047528) consists of 15 blood plasma samples of HIV-positive individuals [27]. For this dataset, we load both DDA and DIA abundance values from the corresponding peptides.txt files, which we received from our analysis of the raw files as described in section C.1. Duplicated peptides are identified by their sequences and grouped together by summing the corresponding abundance values. Matching between DDA and DIA peptides is done via amino acid sequence.

Since for the HEK293-*E.coli* (B1) dataset, the abundance of human proteins is kept constant across all samples, but different amounts of *E. coli* are spiked in, the overall combined abundance of all proteins per sample changes. To make samples comparable, sum normalization of peptide abundance values is performed via summation of the abundance values of human peptides, in the same way as done by Dowell et al. [26]. Normalization is done for both the DDA and the DIA peptide abundance values. No normalization is performed for the HIV blood dataset. However, for both DDA/DIA datasets, we scale the DIA abundance values to have the same mean as the DDA abundance values.

For both DDA/DIA datasets, imputation is done exclusively on the DDA values, using the DIA abundance values only as ground truth for the evaluation of imputation performance. Further, only abundance values missing in the DDA datasets with non-missing corresponding values in the DIA datasets are used for evaluation. Comparison with ground truth DIA abundance values is done differently for both datasets. For the HIV blood dataset, a 1:1 mapping between DDA samples and DIA samples exists, and imputed DDA abundance values of one sample are compared with abundance values of the corresponding DIA sample. For the HEK293-*E. coli* dataset, no such mapping between DDA and DIA samples exists because samples representing technical replicates within one condition group (same amount of spike-in *E. coli* protein digest) cannot be matched between the DDA and DIA datasets. Therefore, ground truth peptide abundance values are computed by taking the median of all non-missing DIA peptide abundance values across technical replicates within a condition group.

##### C.2.3 C: Mixture dataset

We use the dataset from Cox et al. [28] consisting of 60 µg HeLa S3 lysate spiked with either 10 µg or 30 µg *E. coli* K12 lysate resulting in an 1:3 fold change of *E. coli* proteins between the two condition groups of measured replicates. For every ratio, three technical replicates have been created, resulting in a final dataset consisting of six samples and two condition groups. The first condition group consists of the samples called L1, L2, and L3, all spiked with 10 µg of *E. coli* proteins, while the second group consists of the samples called H1, H2, H3, all containing 30 µg of spike-in *E. coli* lysate. Again, we load the dataset from the peptides.txt file output of MaxQuant, which is provided in the corresponding PRIDE repository (PXD000279). Again, we use the "Intensity" columns in that file as our peptide abundance values, while protein-peptide relations are inferred from the "Protein group ID" column. The amino acid sequence of each peptide is again inferred from the "Sequence" column. In addition, sum normalization is done as described above for the HEK293-*E.coli* dataset by normalizing every sample using the sum of peptide abundance values that can be uniquely assigned to human proteins. Further, to use this dataset also for the evaluation of abundance values, we generate a hold-out test set by introducing artificial missing values by masking 10% of non-missing abundance values uniformly at random. We subsequently delete all peptides without a single missing value as described above for the masked datasets.

### D Evaluation procedure for differential expression analysis

For differential expression (DE) discovery for each peptide, the abundance values between biological samples (condition groups) for the HeLa-*E.coli* benchmark dataset (C) are compared using Welch's t-test. P-values are corrected for multiple testing using Benjamini-Hochberg correction.

Ground truth fold changes and the resulting ground truth DEs are inferred from the species-specific mixture rates. All human peptides in the dataset have a ground truth abundance ratio of 1:1 between the two condition groups and form the non-differentially expressed ground truth set, while the *E. coli* peptides show a ground truth abundance ratio of 1:3 between the two condition groups and form the differentially expressed ground truth set. Only peptides that can be uniquely assigned to one species are considered for the evaluation.

### E Implementation

Most of the functionality is implemented in Python (version 3.10.13) using our self-developed, open-source PyProteoNet Python package (`https://github.com/DILiS-lab/pyproteonet`), which we created to foster the development and evaluation of imputation methods for proteomics data. The package provides data structures for the representation of datasets consisting of related proteins and peptides (as well as other molecules) and provides imputation, peptide-to-protein aggregation, and evaluation methods under a unified interface. For a more extensive description of the functionality as well as usage examples, the reader is referred to PyProteoNet's documentation at `https://pyproteonet.readthedocs.io`.

We developed PEPerMINT based on PyProteNet's functionality to transform proteomics datasets into graph structures compatible with the deep graph library (DGL) [29]. Further, we used PyTorch [30] as an automatic differentiation framework and Lightning [31] as a high-level deep learning framework.

For the comparison with other imputation methods, we used the imputation functions provided by PyProteoNet, which in turn wrap common R and Python imputation packages as described in section B. Figures were created using Seaborn [32] and Matplotlib [33].

### F Supplementary evaluation results

Here, we show evaluation results supplementing the results from the main text.

#### F.1 Abundance-based evaluation

Orthogonal to the sample-wise root mean squared error (RMSE) results shown in the main text (Fig. 3A), Fig. S2 shows the mean absolute error computed over all imputed values of a dataset that have corresponding ground truth abundance values. Results match those in the main text, with PEPerMINT outperforming other imputation methods on the majority of datasets.

Fig. S3 shows the RMSE results for the evaluation stratified by the percentage of missing values per peptide for the datasets not shown in Fig. 3B in the main text. As described in the main text, on the majority of datasets, PEPerMINT performance is best out of all evaluated imputation methods independent of the percentage of missing values a peptide shows. However, PEPerMINT usually outperforms other methods by the largest margin on peptides with high percentages of missing values. Error bars in all bar plots (here and in the main text) represent the 95% confidence interval and are generated using the default settings Seaborn's [32] barplot function (bootstrapping, 1000 samples).

Fig. S4 shows the results for statistically comparing the abundance error between imputation methods stratified by the percentage of missing values matching the results shown in Fig. 4 in the main text. Again, it can be seen that PEPerMINT compares favorably against others irrespective of the percentage of missing values per peptide.

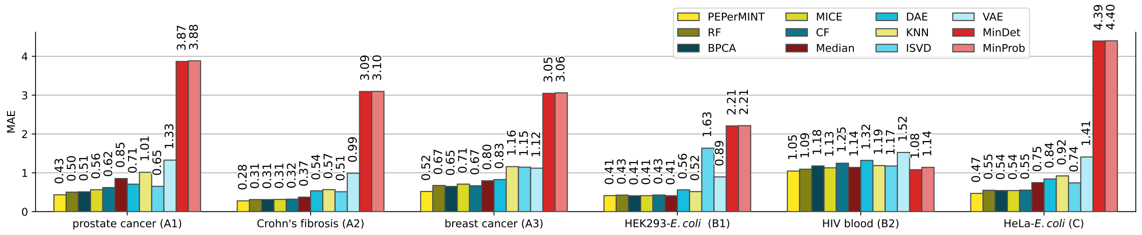

Fig. S2: Dataset-wise imputation performance in terms of MAE comparing our PEPerMINT imputation methods with eleven methods from the literature on six benchmark datasets. As for the RMSE-based evaluation (see Fig. 3A in the main text), PEPerMINT outperforms other imputation methods on the majority of datasets.

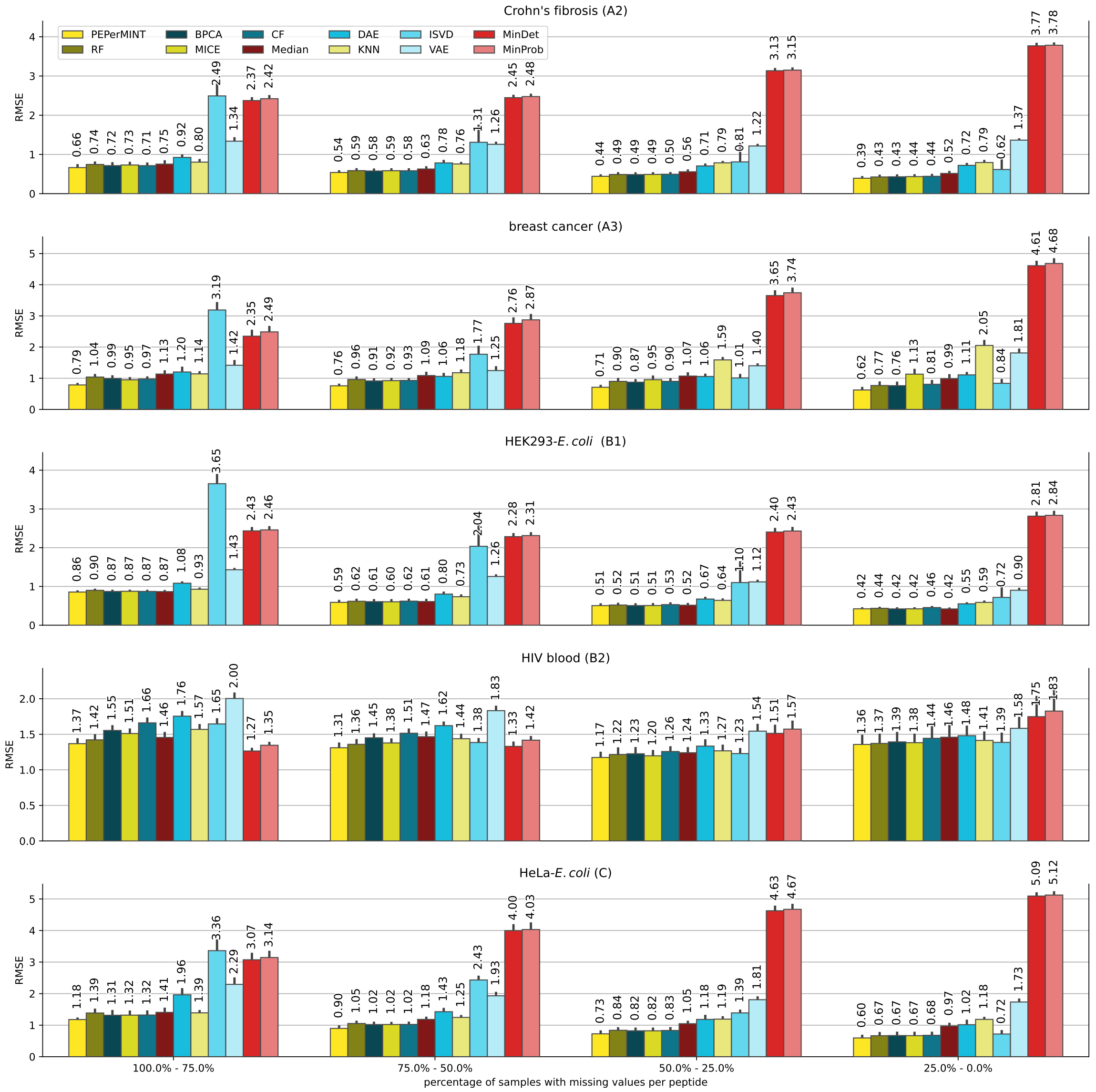

Fig. S3: Sample-wise RMSE per dataset stratified by percentage of missing values per peptide for the five datasets not shown in the main text (see Fig. 3B). Overall, PEPerMINT performs better than other imputation methods independent of the percentage of missing values but has the biggest advantage over other methods on peptides, showing a high percentage of missing values.

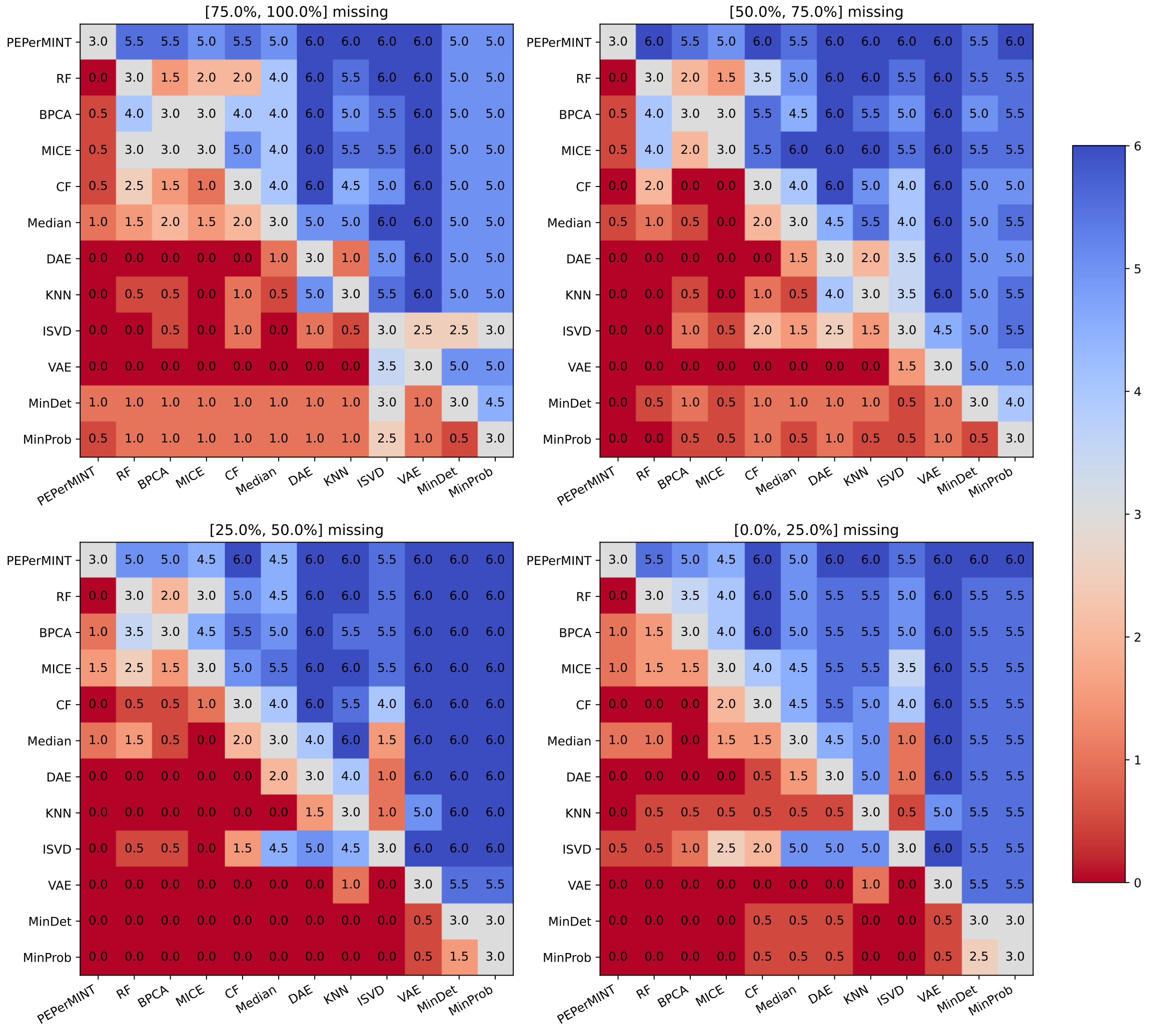

Fig. S4: Pairwise comparison of different imputation methods using a Bonferroni corrected Wilcoxon signed-rank test on the absolute error values stratified by the percentage of missing values per peptide (corresponding to Fig. 4 in the main text). Each cell value counts on how many datasets the imputation method given by the row performs significantly better than the method given by the column. If none of the two compared methods performs significantly better, they both get a count of 0.5. PEPerMINT outperforms the other methods on the majority of datasets, irrespective of the percentage of missing values per peptide.

#### F.2 Evaluation of differential expression prediction

Fig. S5 shows the precision-recall curve for the DE analysis with 5% thresholds marked with dots. Similar to the ROC curve shown in Fig. 5 in the main text, PEPerMINT has the highest area under the curve.

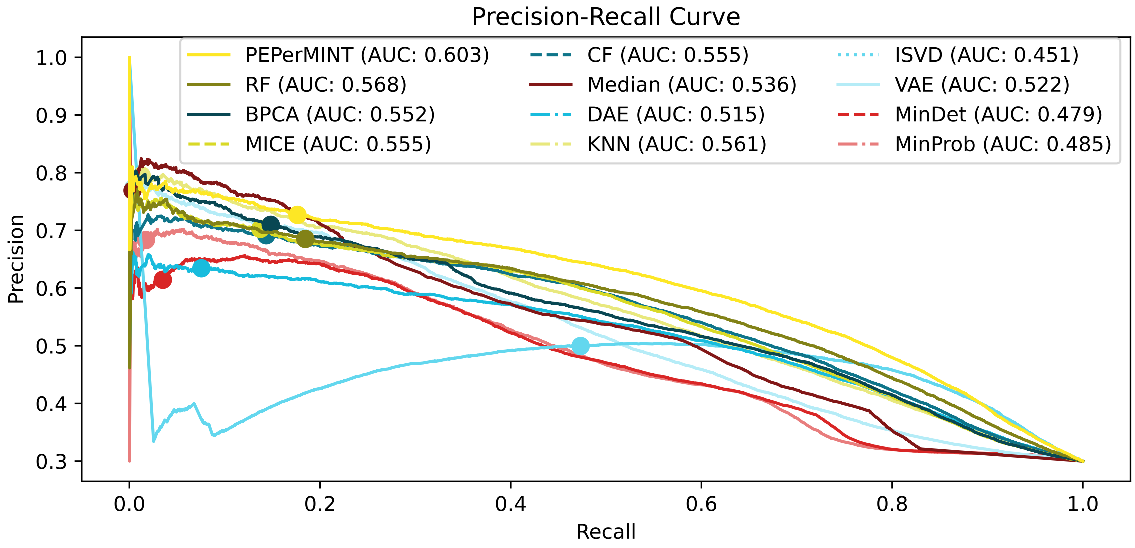

Fig. S5: Precision recall curve corresponding to Fig. 5 in the main text and visualizing the performance of predicting differentially expressed peptides on the MaxLFQ benchmark dataset imputed with different imputation methods with 5% FDR thresholds marked by dots. Our PEPerMINT imputation method has the largest area under the curve (AUC).

#### F.3 Predicted uncertainty of imputed values

Fig. S6 visualizes the estimated uncertainty for imputed values for the datasets not shown in Fig. 6 in the main text. As with the figures in the main text, it can be seen that imputed values with small absolute errors with respect to ground truth abundance values tend to have lower predicted uncertainty values assigned. It can also be seen that high abundance values tend to show lower uncertainties.

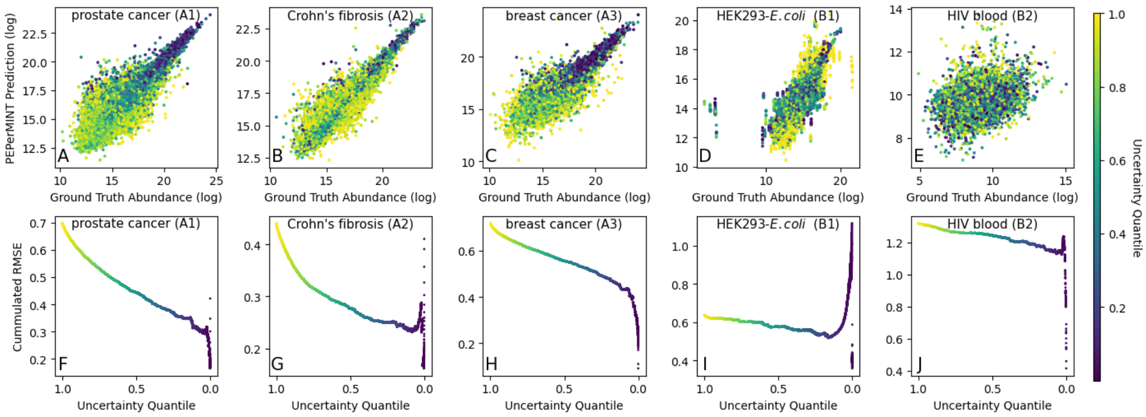

Fig. S6: Visualization of uncertainty predicted for the imputed values by our PEPerMINT imputation method for the datasets not shown in Fig. 6 in the main text. For each dataset, two plots are shown. A-E: Scatter plot showing the imputed abundance values against their corresponding ground truth values with different uncertainties visible as different colors (blue: low, yellow: high). It can be seen that imputed values with small absolute errors with respect to ground truth abundance values tend to have lower predicted uncertainty values assigned. It can also be seen that high abundance values tend to show lower uncertainties. F-J: The imputation error when ordering imputed values by their predicted uncertainty and computing the MSE over iteratively grown uncertainty quantiles. It can be seen that only considering imputed values showing a low uncertainty also results in a lower MSE. One exception is the HEK293-*E.coli* dataset, which contains several outliers with very low ground truth abundance values (see scatter plot D) explaining the different results.

#### F.4 Impact of predicted uncertainty on differential expression analysis

Fig. S7 shows ROC curves stratified by predicted uncertainty for PEPerMINT. The same curves zoomed in to lower FDR rates but without the curves showing DE analysis performance without imputation and DE analysis only on peptides without any missing values are shown in Fig. 6C in the main text. For each curve, only peptides with all their abundance values in all six samples, either non-missing or imputed with an uncertainty below a certain threshold, are evaluated (resulting in different amounts of evaluated peptides). Overall, it can be seen that restricting evaluation to peptides with lower uncertainty also results in a higher AUC. Interestingly, it can also be seen that including imputed values with low uncertainty ($\leq 0.2$ uncertainty quantile, $\leq 0.4$ uncertainty quantile) can actually achieve a better performance than only considering peptides without any missing values (fully measured only), both in terms of AUC as well as in the amount of correctly identified differentially expressed peptides at 5% FDR level. One reason might be that low uncertainties only get assigned to imputed values, where the remaining non-missing values across samples are consistent and allow a clear categorization into differentially expressed and non-differentially expressed. Thus, adding imputed values in the order of their predicted uncertainty by PEPerMINT up to intermediate thresholds (here: 0.4) shows improved TPRs for DE while also gaining insights on 33500 instead of only 18100 peptides without imputation. Thus, a decisive gain in downstream analysis is achieved by our PEPerMINT-based approach.

In addition, the plot shows results for DE analysis without imputation. In that case, missing values are filtered out before performing Welch's t-test. P-values for peptides where the test cannot be applied (e.g., because of insufficient non-missing abundance values) are set to 1.0. Performance in DE analysis is the lowest for this approach.

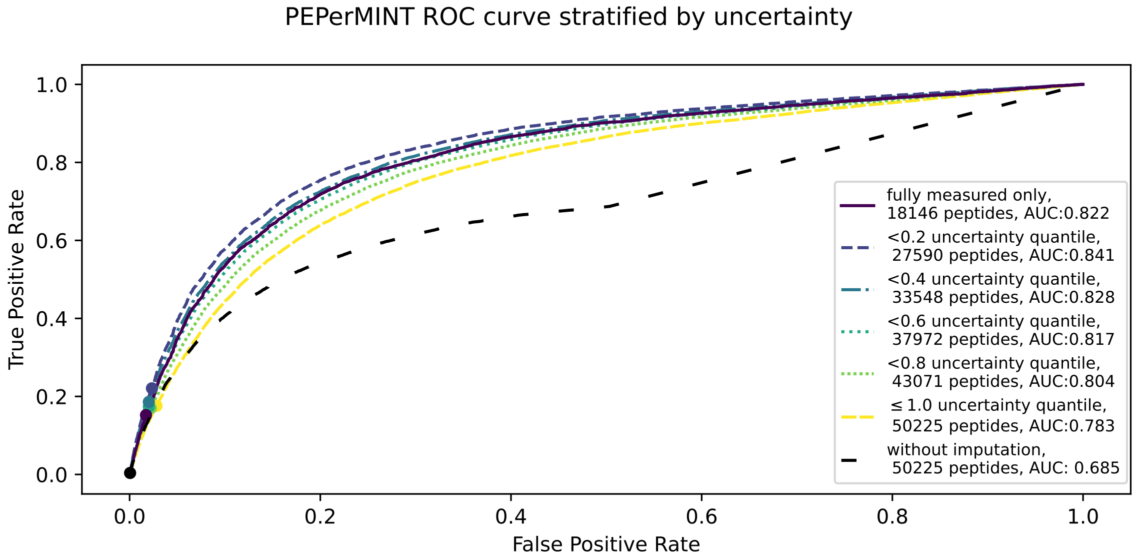

Fig. S7: Full ROC curves for the ROC curves partly shown in Fig. 6C in the main text showing DE analysis performance when stratified by uncertainty. 5% FDR thresholds are marked by dots

### G Runtime

For each imputation method, we report the runtime on every evaluated dataset (see Fig. S8). For our PEPerMINT method, we only consider the runtime of the imputation run on a specific dataset without the generation of peptide sequence embeddings, as those can be precomputed and cached (as done by our implementation). While runtimes differ by orders of magnitude between methods, it can be observed that well-performing and more complex methods like PEPerMINT, BPCA, or RF also have the largest runtime. The runtime of our PEPerMINT imputation method is roughly similar to that of BPCA imputation. However, it should be noted that exact runtimes can differ with respect to the hardware used. For example, our PEPerMINT imputation, as well as both autoencoder imputation methods (VAE, DAE), were executed on a GPU (NVIDIA T4), while all other imputation methods were run on a CPU (AMD EPYC 7742).

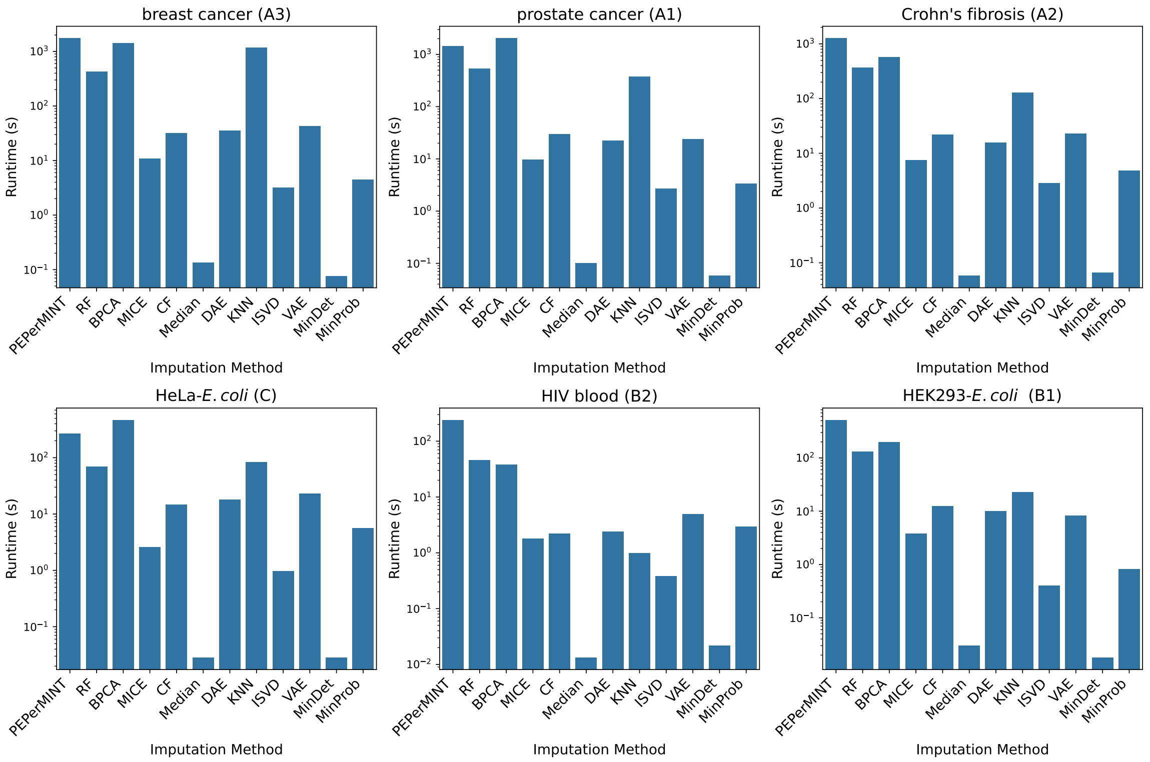

Fig. S8: Runtime of all imputation methods on the different datasets.

### H Ablation study

To assess how different key components of PEPerMINT (see Fig. S1) contribute to its performance, we performed an ablation study focusing on the following aspects:

- Role of the peptide graph: To assess the role of the peptide graph, we replaced all edges with random edges.
- Role of the peptide sequence embeddings: To assess the role of sequence embeddings, we replaced every embedding with a zero vector.
- Role of the skip connection bypassing the graph attention layer: The skip connections bypass the graph attention layer to combine both local information of a peptide (from the skip connection) and information from its neighbors (from the graph attention layer). To assess their role, we remove them from our neural network architecture. We adapt the subsequent layers to accommodate the reduced size of the latent representation.

We restrict the ablation study to our masked prostate cancer dataset. Fig S9 shows the sample-wise RMSE and MAE results for our complete PEPerMINT model as well as for the different ablation configurations described above. All of PEPerMINT's components seem to have a positive effect on imputation performance, as can be seen by the reduced performance when they are removed. Interestingly, the skip connection has the largest effect on the performance. One explanation might be that the abundance of a particular peptide node cannot be solely inferred by aggregating information from neighboring peptides without taking the information of the particular peptide node into account. This relates to the problem of oversmoothing and its mitigation via skip connections as it is commonly reported for graph neural networks [34].

The observations stated above are further confirmed when stratifying the ablation study results by the fraction of missing values per peptide as shown in Fig. S10. It can be seen that sequence embeddings and connections between peptides of the same protein, as given by the peptide graph, seem to be most helpful for peptides with high fractions of missing values across samples. In contrast, the positive effect of the skip connection bypassing the graph attention layer seems to be most pronounced on peptides with a low fraction of missing values. This matches well with our intuition that additional information like those provided by sequence embeddings and the peptide graph are most helpful for peptides with little abundance information (high fraction of missing values). However, single missing values of peptides that have non-missing abundance values in many other samples can often be inferred from those non-missing values of other samples (achieved by the skip connection in PEPerMINT's architecture).

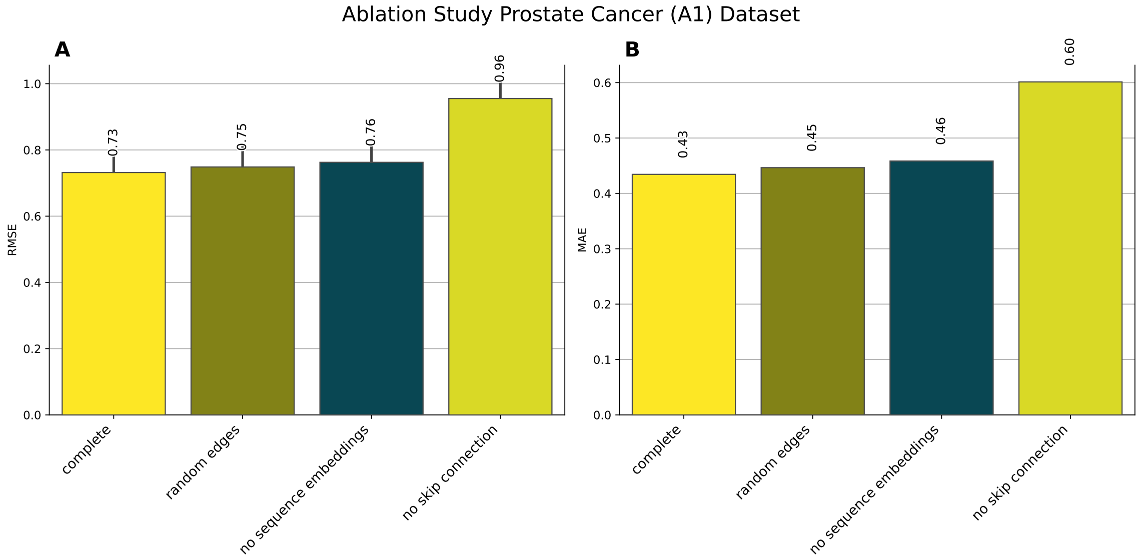

Fig. S9: Ablation study results with respect to sample-wise RMSE (A) and MAE (B). Bars visualize PEPerMINT's performance with different components disabled compared with the performance of the full model (complete). All of PEPerMINT's components seem to have a positive effect on performance, as can be seen by the reduced performance when they are removed. The skip connection bypassing the graph attention layer seems to be the most important.

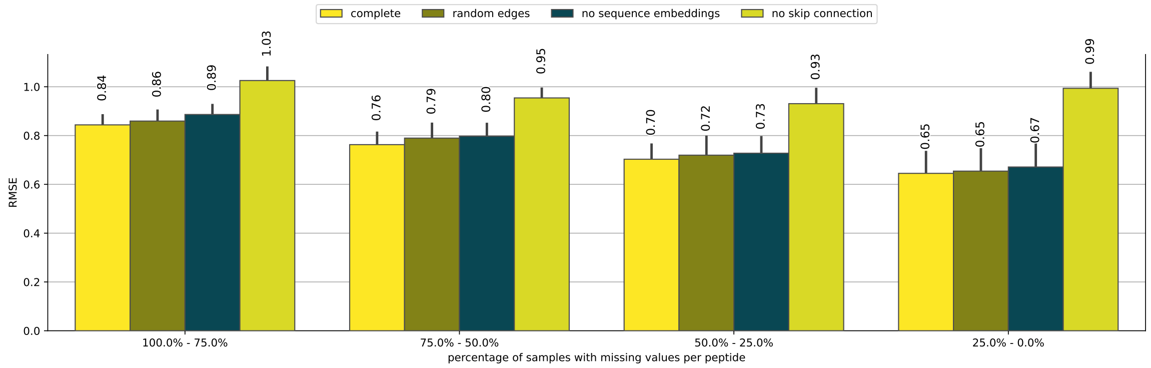

Fig. S10: Ablation study results with respect to sample-wise RMSE stratified by fraction of missing values per peptide. Bars visualize PEPerMINT's performance with different components disabled compared with the performance of the full model (complete). Sequence embeddings and peptide graph edges seem to be most helpful for peptides with high fractions of missing values across samples. In contrast, the positive effect of the skip connection is most pronounced on peptides with a low fraction of missing values.
